# *Pdgfra* and *Pdgfrb* genetically interact in the murine neural crest cell lineage to regulate migration and proliferation

**DOI:** 10.1101/2020.07.29.227306

**Authors:** Julia Mo, Robert Long, Katherine A. Fantauzzo

**Affiliations:** Department of Craniofacial Biology, School of Dental Medicine, University of Colorado Anschutz Medical Campus, Aurora, CO, United States of America

**Keywords:** Pdgfra, Pdgfrb, neural crest, craniofacial, migration, proliferation

## Abstract

Cranial neural crest cells (cNCCs) are migratory, multipotent cells that originate from the forebrain to the hindbrain and eventually give rise to the bone and cartilage of the frontonasal skeleton, among other derivatives. Signaling through the two members of the platelet-derived growth factor receptor (PDGFR) family of receptor tyrosine kinases, alpha and beta, plays critical roles in the cNCC lineage to regulate craniofacial development during murine embryogenesis. Further, the PDGFRs have been shown to genetically interact during murine craniofacial development at mid-to-late gestation. Here, we examined the effect of ablating both *Pdgfra* and *Pdgfrb* in the murine NCC lineage on earlier craniofacial development and determined the cellular mechanisms by which the observed phenotypes arose. Our results confirm a genetic interaction between the two receptors in this lineage, as phenotypes observed in an allelic series of mutant embryos often worsened with the addition of conditional alleles. The defects observed here were shown to stem from reduced cNCC stream size and aberrant cNCC directional migration, as well as decreased proliferation of the facial mesenchyme upon combined decreases in PDGFRα and PDGFRβ signaling. Importantly, we found that PDGFRα plays a predominant role in cNCC migration whereas PDGFRβ primarily contributes to proliferation of the facial mesenchyme. Our findings provide insight into the distinct mechanisms by which PDGFRα and PDGFRβ signaling regulate cNCC activity and subsequent craniofacial development in the mouse embryo.

## Introduction

Neural crest cells (NCCs) are migratory, multipotent cells that play critical roles in vertebrate development. NCCs arise at the border of the neural ectoderm, undergo an epithelial to mesenchymal transition and subsequently delaminate from the cranial neural folds or dorsal neural tube during mammalian embryogenesis. Cranial NCCs (cNCCs) originate from the forebrain to the hindbrain and eventually give rise to the bone and cartilage of the frontonasal skeleton, as well as the cartilages of the jaw, middle ear, hyoid and thyroid, among other derivatives (Trainor, 2005; Mayor and Theveneau, 2013). At approximately embryonic day (E) 9.5 in the mouse, the process of craniofacial development begins with the formation of five facial prominences populated by post-migratory cNCCs. These prominences include the frontonasal prominence, a pair of maxillary prominences and a pair of mandibular prominences. Subsequent formation of the nasal pits divides the frontonasal prominence into the medial and lateral nasal processes, which will eventually fuse and give rise to the nostrils. A second fusion event occurs between the medial nasal processes and the maxillary prominences to form the upper lip. At this time the secondary palatal shelves first appear as morphologically distinct outgrowths on the oral side of the maxillary prominences. The shelves grow downward as they extend from the maxillae such that they are positioned on either side of the tongue. The palatal shelves elevate to a horizontal, apposing position above the tongue with development of the jaw and grow towards the midline. The palatal shelves fuse with one another upon meeting and eventually with two derivatives of the medial nasal processes, the primary palate anteriorly and the nasal septum superiorly, resulting in a continuous palate that separates the oral and nasal cavities (Bush and Jiang, 2012). The complex morphogenetic process of craniofacial development requires a precise interplay of multiple cell and tissue types. As such, defects in craniofacial development, including cleft lip and palate, comprise one of the most prevalent birth defects in humans (Parker et al., 2010).

Signaling through the platelet-derived growth factor receptor (PDGFR) family of receptor tyrosine kinases plays a critical role in human craniofacial development. There are four PDGF ligands in mammals, PDGF-A-D, which signal through two receptors, PDGFRa and PDGFRβ. The homodimers PDGF-AA and PDGF-CC have been shown to solely activate PDGFRa signaling *in vivo* during mammalian development (Boström et al., 1996; Soriano, 1997; Ding et al., 2004), while PDGF-BB exclusively activates PDGFRβ signaling (Levéen et al., 1994; Soriano, 1994). Ligand binding induces PDGFR dimerization and activation of cytoplasmic tyrosine kinase domains, which in turn autophosphorylate intracellular tyrosine residues. Signaling molecules bind to specific phosphorylated residues in the cytoplasmic domains of the receptors and mediate downstream cellular responses through various intracellular signaling pathways (Heldin and Westermark, 1999). Heterozygous missense mutations in the human *PDGFRA* coding region and single base-pair substitutions in the 3’ untranslated region are associated with nonsyndromic cleft palate (Rattanasopha et al., 2012). Further, single-nucleotide polymorphisms in the regulatory region of *PDGFC* which repress transcriptional activity of the promoter are associated with cleft lip and palate (Choi et al., 2009). Alternatively, heterozygous missense mutations in *PDGFRB* have been shown to cause Kosaki overgrowth syndrome (OMIM 616592) and Penttinen syndrome (OMM 601812), both of which are characterized by facial dysmorphism, among other defects (Johnston et al., 2015; Takenouchi et al., 2015).

The roles of PDGFRα and PDGFRβ in human craniofacial development are evolutionarily conserved in the mouse. Targeted disruption of *Pdgfra* in mice results in embryonic lethality during mid-gestation, with homozygous null embryos exhibiting facial clefting, subepidermal blebbing, edema, hemorrhaging, cardiac outflow tract defects, abnormalities in neural tube development, abnormally patterned somites and extensive skeletal defects affecting cNCC derivatives in the frontonasal skeleton, as well as non-NCC-derived axial skeletal elements (Soriano, 1997). These defects are phenocopied in embryos lacking both *Pdgfa* and *Pdgfc* (Ding et al., 2004). *Pdgfra* is expressed in migrating cNCCs and in the cNCC-derived mesenchyme of the facial processes during mid-gestation, among other sites, while its ligands, *Pdgfa* and *Pdgfc*, are reciprocally expressed in the overlying epithelium (Morrison-Graham et al., 1992; Orr-Urtreger and Lonai, 1992; Ding et al., 2000; Hamilton et al., 2003; He and Soriano, 2013; Fantauzzo and Soriano, 2016). Conditional ablation of *Pdgfra* in the NCC lineage using the *Wnt1-Cre* driver (Danielian et al., 1998) generates a subset of the null phenotypes, including facial clefting, midline hemorrhaging, aortic arch defects and thymus hypoplasia (Tallquist and Soriano, 2003; He and Soriano, 2013). *Pdgfra^fl/fl^;Wnt1-Cre^+/Tg^* embryos exhibit a delay in NCC migration into the frontonasal prominence at E9.5 and fewer NCCs in pharyngeal arches 3-6 at E10.5, with bifurcation of the streams entering these arches in a subset of embryos (He and Soriano, 2013). Additionally, these embryos have decreased proliferation in the frontonasal and medial nasal processes at E9.5 and E11.5, respectively (He and Soriano, 2013). Similarly, PDGFRα signaling has been shown to regulate cell survival and proliferation of the cNCC-derived mesenchyme contributing to the palatal shelves at E13.5 (Fantauzzo and Soriano, 2014). Conditional ablation of *Pdgfra* specifically in cNCCs using the *Sox10ER^T2^CreER^T2^* driver and following administration of tamoxifen at E7.5 likewise leads to fewer NCCs in the craniofacial region at E10.5, decreased proliferation in the medial nasal process at E11.5 and eventual frontonasal dysplasia (He and Soriano, 2015). Interestingly, use of this driver revealed a novel requirement for PDGFRα in the mandible, as *Pdgfra^fl/fl^;Sox10ER^T2^CreER^T2^* embryos additionally exhibited decreased proliferation in the mandibular mesenchyme at E11.5 and mandibular hypoplasia at E16.5 (He and Soriano, 2015). Conversely, both *Pdgfrb-* and *Pdgfb-deficient* mice die perinatally and exhibit edema, hemorrhaging, cardiac ventricular septal defects, thrombocytopenia, anemia and kidney defects (Levéen et al., 1994; Soriano, 1994). *Pdgfrb* is also expressed in the embryonic craniofacial mesenchyme (Soriano, 1994; Fantauzzo and Soriano, 2016; McCarthy et al., 2016) and ablation of *Pdgfrb* in the NCC lineage results in increased nasal septum width, delayed palatal shelf development and subepidermal blebbing in a subset of embryos (Fantauzzo and Soriano, 2016). Though the etiology of these defects is currently unknown, *Pdgfrb^fl/fl^;Wnt1-Cre^+/Tg^* embryos do not have obvious defects in NCC migration into the facial processes and pharyngeal arches at E8.5-E10.5 (Fantauzzo and Soriano, 2016).

The PDGFRs have been shown to genetically interact during murine craniofacial and heart development. While a previous skeletal analysis in which both *Pdgfra* and *Pdgfrb* were simultaneously conditionally ablated in the NCC lineage did not detect additional frontonasal midline defects in double-homozygous mutant embryos beyond those observed in *Pdgfra^fl/fl^;Wnt1-Cre^+/Tg^* embryos (McCarthy et al., 2016), malformations in the basisphenoid, alisphenoid and hyoid bones at E17.5, as well as defects in multiple cardiac NCC derivatives at E14.5-E18.5, were observed that were more severe than those found in either single-homozygous mutant alone (Richarte et al., 2007; McCarthy et al., 2016). The latter phenotype was shown to arise from cardiac NCC migration defects into the outflow tract as early as E10.5 and not from defects in proliferation nor survival of cells in the conotruncal region between E10.5-E12.5 (Richarte et al., 2007). Phosphatidylinositol 3-kinase (PI3K) has been identified as the main downstream effector of PDGFRα signaling during murine embryonic development (Klinghoffer et al., 2002). Embryos homozygous for an autophosphorylation mutant knock-in allele (*Pdgfra^PI3K^*) in which PDGFRα is unable to bind PI3K die perinatally and display a cleft palate, among other defects (Klinghoffer et al., 2002; Fantauzzo and Soriano, 2014), which is less severe than the complete facial clefting phenotype observed in *Pdgfra^fl/fl^;Wnt1-Cre^+/Tg^* embryos (Tallquist and Soriano, 2003; He and Soriano, 2013). While *Pdgfra^PI3K/PI3K^* embryos do not exhibit NCC migration defects at E9.5-E10.5 (He and Soriano, 2013), primary mouse embryonic palatal mesenchyme cells (MEPMs) derived from E13.5 *Pdgfra^PI3K/PI3K^* embryos fail to proliferate in response to PDGF-AA ligand treatment (He and Soriano, 2013; Fantauzzo and Soriano, 2014). When the constitutive *Pdgfra^PI3K^* allele was combined with the conditional *Pdgfrb^fl^* allele and the *Wnt1-Cre* driver, E13.5 double-homozygous mutant embryos had an overt facial clefting phenotype not observed in either single-homozygous mutant (Fantauzzo and Soriano, 2016). Further, introduction of a single *Pdgfrb^fl^* allele exacerbated the midline defects observed in *Pdgfra^PI3K/PI3K^* skeletons at E16.5 such that *Pdgfra^PI3K/PI3K^;Pdgfrb^+/fl^;Wnt1-Cre^+/Tg^* skeletons additionally exhibited upturned and clefted nasal cartilage, a widening of the gap between the premaxilla bones and generalized broadening of the skull (Fantauzzo and Soriano, 2016), similar to the craniofacial skeletal defects observed upon conditional ablation of *Pdgfra* in the NCC lineage (Tallquist and Soriano, 2003; He and Soriano, 2013).

To examine the effect of ablating both *Pdgfra* and *Pdgfrb* in the murine NCC lineage on earlier craniofacial development and to determine the cellular mechanisms by which the observed phenotypes arise, we analyzed an allelic series of mutant embryos. Our results confirm a genetic interaction between the two receptors in this lineage and demonstrate that PDGFRα plays a predominant role in cNCC migration whereas PDGFRβ exerts its effect primarily through the regulation of proliferation in the facial mesenchyme.

## Materials and Methods

### Mouse strains

All animal experimentation was approved by the Institutional Animal Care and Use Committee of the University of Colorado Anschutz Medical Campus. *Pdgfra^tm8Sor^* mice (Tallquist and Soriano, 2003), referred to in the text as *Pdgfra^fl^; Pdgfrb^tm11Sor^* mice (Schmahl et al., 2008), referred to in the text as *Pdgfrb^fl^; H2afv^Tg(Wnt1-cre)11Rth^* mice (Danielian et al., 1998), referred to in the text as *Wnt1-Cre^Tg^*; and *Gt(ROSA)26Sor^tm4(ACTB-tdTomato,-EGFP)Luo^* mice (Muzumdar et al., 2007), referred to in the text as *ROSA26^mTmG^*, were maintained on a 129S4 coisogenic genetic background. Statistical analyses of Mendelian inheritance were performed with the GraphPad QuickCalcs data analysis resource (GraphPad Software, Inc., La Jolla, CA, USA) using a chi-square test. Statistical analyses of litter sizes were performed Prism 8 (GraphPad Software, Inc.) using a two-tailed, unpaired t-test with Welch’s correction.

### Morphological analysis

Embryos were dissected at multiple timepoints (day of plug considered 0.5 days) in 1x phosphate buffered saline (PBS) and fixed overnight at 4°C in 4% paraformaldehyde (PFA) in PBS. Embryos were photographed using an Axiocam 105 color digital camera (Carl Zeiss, Inc., Thornwood, NY, USA) fitted onto a Stemi 508 stereo microscope (Carl Zeiss, Inc.). Distances between nasal pits were measured using Photoshop software v 21.1.1 (Adobe, San Jose, CA, USA). Statistical analyses were performed with Prism 8 (GraphPad Software, Inc.) using a two-tailed, unpaired t-test with Welch’s correction and Welch and Brown-Forsythe ANOVA tests.

### Whole-mount DAPI staining

Whole-mount 4’,6-diamidino-2-phenylindole (DAPI) staining was performed according to a previously published protocol (Sandell et al., 2012), with the exception that staining was performed with 10 μg/mL DAPI (Sigma-Aldrich Corp., St. Louis, MO, USA) for 1 hr at room temperature. Embryos were photographed using an Axiocam 506 mono digital camera (Carl Zeiss, Inc.) fitted onto an Axio Observer 7 fluorescence microscope (Carl Zeiss, Inc.). Extended Depth of Focus was applied to z-stacks using ZEN Blue software (Carl Zeiss, Inc.) to generate images with the maximum depth of field. An Unsharp Mask was applied to select images of NCC streams at E10.5 using ImageJ software (version 2.0.0-rc-69/1.52p; National Institutes of Health) with radius 40 pixels and mask weight 0.90. Anterior-posterior heights and dorsal-ventral lengths of NCC streams in at least three embryos per genotype per timepoint were measured using ZEN Blue software (Carl Zeiss, Inc.). Statistical analyses were performed with Prism 8 (GraphPad Software, Inc.) using a two-tailed, unpaired t-test with Welch’s correction and Welch and Brown-Forsythe ANOVA tests.

### TUNEL assay

Embryos were fixed in 4% PFA in PBS and infiltrated with 30% sucrose in PBS before being mounted in O.C.T. compound (Sakura Finetek USA Inc., Torrance, CA, USA). Sections (8 μm) were deposited on glass slides. Apoptotic cells were identified using the *In Situ* Cell Death Detection Kit, Fluorescein (Sigma-Aldrich Corp.) according to the manufacturer’s instructions for the treatment of cryopreserved tissue sections. Sections were mounted in VECTASHIELD^®^ Antifade Mounting Medium with DAPI (Vector Laboratories, Burlingame, CA, USA) and photographed using an Axiocam 506 mono digital camera (Carl Zeiss, Inc.) fitted onto an Axio Observed 7 fluorescence microscope (Carl Zeiss, Inc.). All positive signals were confirmed by DAPI staining. The percentage of TUNEL-positive cells was determined in three embryos per genotype per timepoint. Statistical analyses were performed with Prism 8 (GraphPad Software, Inc.) using a two-tailed, unpaired t-test with Welch’s correction and Welch and Brown-Forsythe ANOVA tests.

### Ki67 Immunofluorescence analysis

Sections (8 μm) of PFA-fixed, sucrose-infiltrated, O.C.T-mounted embryos were deposited on glass slides. Sections were fixed in 4% PFA in PBS with 0.1% Triton X-100 for 10 min and washed in PBS with 0.1% Triton-X 100. Sections were blocked for 1 hr in 5% normal donkey serum (Jackson ImmunoResearch Inc., West Grove, PA, USA) in PBS and incubated overnight at 4°C in anti-Ki67 primary antibody (1:300; Invitrogen, Carlsbad, CA, USA) in 1% normal donkey serum in PBS. After washing in PBS, sections were incubated in Alexa Fluor 488-conjugated donkey anti-rabbit secondary antibody (1:1,000; Invitrogen) diluted in 1% normal donkey serum in PBS with 2 μg/mL DAPI (Sigma-Aldrich Corp.) for 1 hr. Sections were mounted in Aqua Poly/Mount mounting medium (Polysciences, Inc., Warrington, PA, USA) and photographed using an Axiocam 506 mono digital camera (Carl Zeiss, Inc.) fitted onto an Axio Observer 7 fluorescence microscope (Carl Zeiss, Inc.). All positive signals were confirmed by DAPI staining. The percentage of Ki67-positive cells was determined in three embryos per genotype per timepoint. Statistical analyses were performed with Prism 8 (GraphPad Software, Inc.) using a two-tailed, unpaired t-test with Welch’s correction and Welch and Brown-Forsythe ANOVA tests.

### Cell culture and growth assays

Primary mouse embryonic palatal mesenchyme (MEPM) cells were isolated from the palatal shelves of embryos dissected at E13.5 in PBS and cultured in medium (Dulbecco’s modified Eagle’s medium (GIBCO, Invitrogen) supplemented with 50 U/mL penicillin (GIBCO), 50 μg/mL streptomycin (GIBCO) and 2 mM L-glutamine (GIBCO)) containing 10% fetal bovine serum (FBS; HyClone Laboratories, Inc., Logan, UT, USA) as previously described (Bush and Soriano, 2010). For cell growth assays, 11,500 passage 2 MEPM cells were seeded into wells of a 24-well plate and cultured in medium containing 10% FBS. After 24 hrs, medium was aspirated and replaced with fresh medium containing 10% FBS (growth medium) or 0.1% FBS (starvation medium). After an additional 24 hrs, select wells were treated daily with 10 ng/mL PDGF-AA, PDGF-BB or PDGF-DD ligand (R&D Systems, Minneapolis, MN, USA) for up to 4 d. Cells were subsequently fixed in 4% PFA in PBS, stained with 0.1% crystal violet in 10% ethanol, extracted with 10% acetic acid and the absorbance measured at 590 nm. Data represent results from three independent trials, each consisting of MEPMs derived from one heterozygous embryo and at least one conditional knock-out littermate. Statistical analyses were performed with Prism 8 (GraphPad Software, Inc.) using a two-tailed, unpaired t-test with Welch’s correction and Welch and Brown-Forsythe ANOVA tests.

## Results

### Pdgfra *and* Pdgfrb *genetically interact in the NCC lineage*

To examine the effect of ablating both *Pdgfra* and *Pdgfrb* in the NCC lineage on mid-gestation craniofacial development, we intercrossed *Pdgfra^fl/fl^;Pdgfrb^fl/fl^* mice with *Pdgfra^+/fl^;Pdgfrb^+/fl^;Wnt1-Cre^+/Tg^* mice and harvested the resulting progeny at E10.5 for gross morphological examination. Double-homozygous mutant embryos were recovered at Mendelian frequencies at this timepoint (16 embryos vs. 14 expected embryos out of 109 total, χ^2^ two-tailed p = 0.4915) (Table 1). A small percentage of embryos across several of the eight allele combinations from the intercrosses exhibited an abnormal head shape due to a misshapen forebrain and/or midbrain, blebbing of the surface ectoderm in the facial region and/or facial hemorrhaging (Table 1). Further, 18% of *Pdgfra^fl/fl^;Pdgfrb^+/fl^;Wnt1-Cre^+/Tg^* embryos displayed ventral body wall closure defects (n = 11) (Table 1).

**Table 1.**
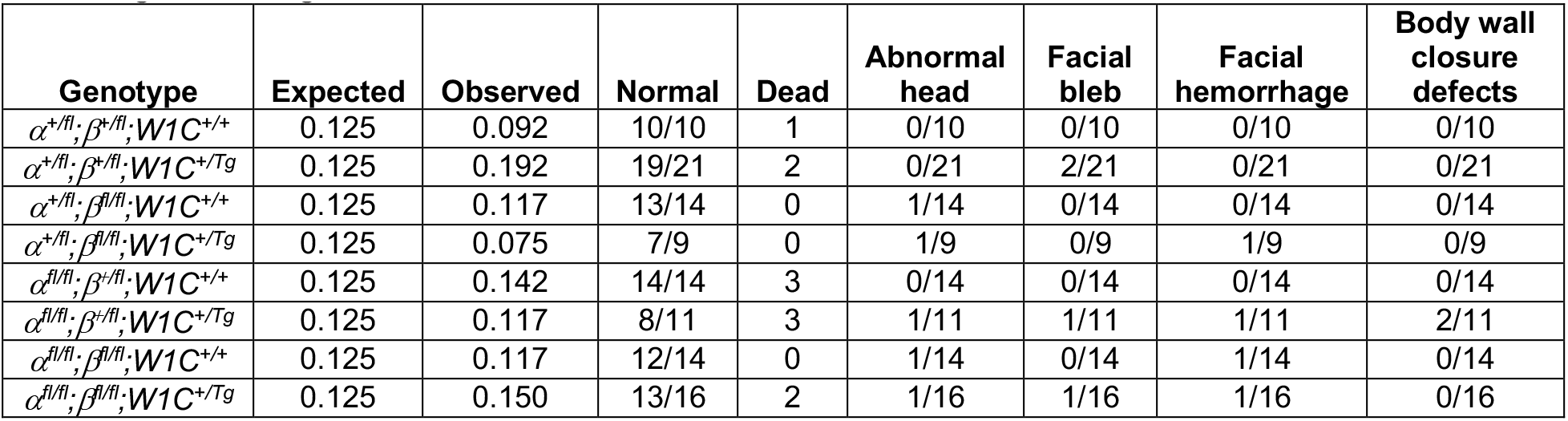
Phenotypes of E10.5 embryos from intercrosses of *Pdgfra^fl/fl^;Pdgfrb^fl/fl^* mice with *Pdgfra^+/fl^;Pdgfrb^+/fl^;Wnt1-Cre^+/Tg^* mice.

We next measured the distance between nasal pits in E10.5 embryos as a readout of defects at the facial midline, revealing a significant difference in measurements across one control (*Pdgfra^+/fl^;Pdgfrb^+/fl^;Wnt1-Cre^+/+^*) and the four experimental genotypes containing the *Wnt1-Cre* transgene (Welch’s ANOVA test p = 0.0001; Brown-Forsythe ANOVA test p < 0.0001). The distance between nasal pits was significantly increased in *Pdgfra^+/fl^;Pdgfrb^fl/fl^;Wnt1-Cre^+/Tg^* embryos (1246 ± 34.12 μm, p = 0.0304), *Pdgfra^fl/fl^;Pdgfrb^+/fl^;Wnt1-Cre^+/Tg^* embryos (1430 ± 31.08 μm, p < 0.0001) and double-homozygous mutant embryos (1349 ± 22.44 μm, p = 0.0006) compared to control *Pdgfra^+/fl^;Pdgfrb^+/fl^;Wnt1-Cre^+/+^* embryos (1107 ± 46.41 μm) (Figure 1). While double-heterozygous mutant embryos had a larger distance between nasal pits than control embryos, this difference was not statistically significant (Figure 1). Interestingly, the greatest distance between nasal pits was observed in *Pdgfra^fl/fl^;Pdgfrb^+/fl^;Wnt1-Cre^+/Tg^* embryos, though this distance was not significantly different between these and double-homozygous mutant embryos (Figure 1).

**Figure 1.**
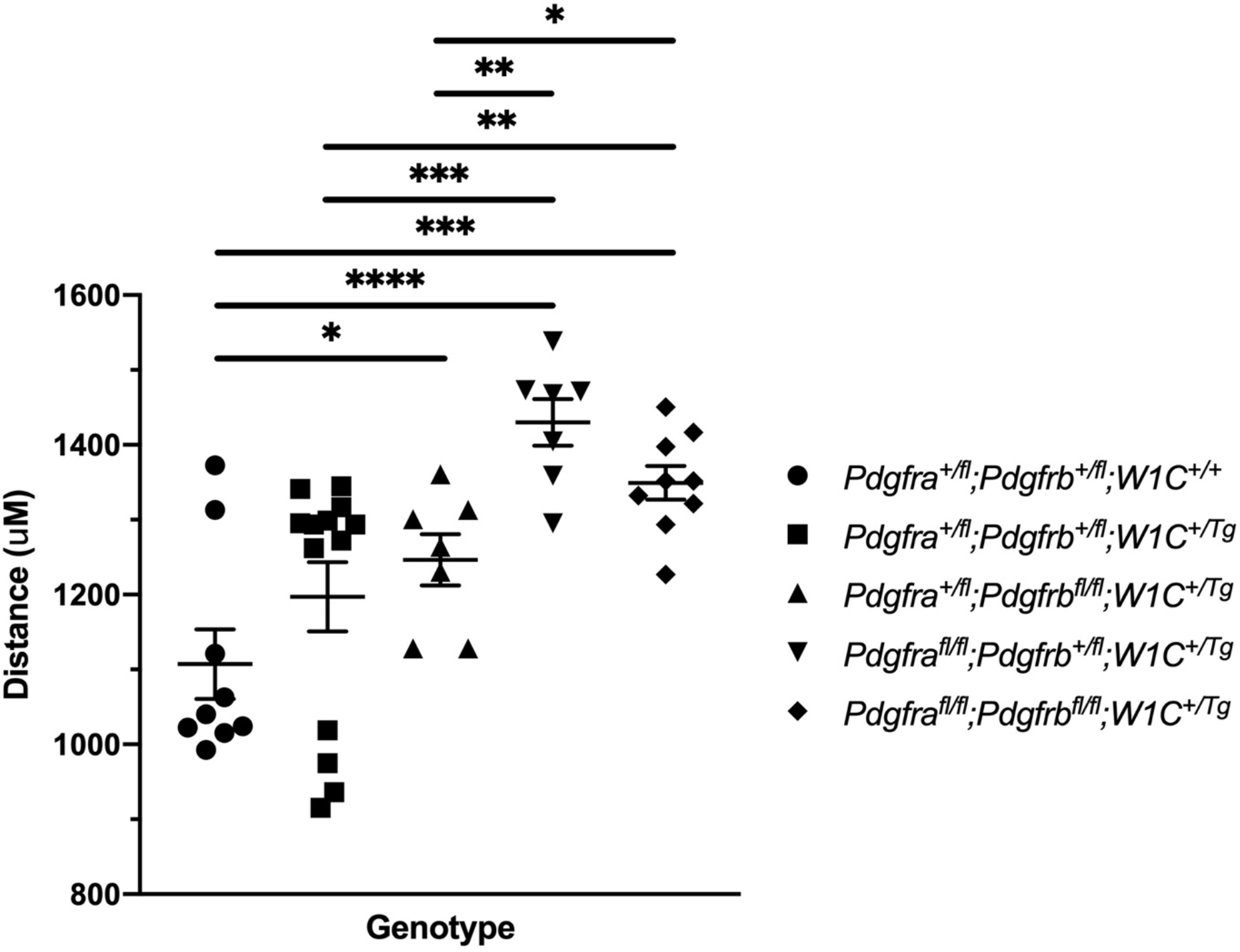
Ablation of *Pdgfra* and *Pdgfrb* in the NCC lineage leads to increased distances between the nasal pits at mid-gestation. Scatter dot plot depicting the distance (μm) between nasal pits across five genotypes at E10.5. Data are presented as mean ± SEM. *, p < 0.05; **, p < 0.01; ***, p < 0.001; ****, p < 0.0001.

To determine whether the above craniofacial phenotypes persisted or worsened at later timepoints, embryos were harvested at E13.5 from the same intercrosses. While the presence of the *Wnt1-Cre* transgene always exacerbated E13.5 facial phenotypes, facial blebbing was detected in a subset of embryos upon combination of at least three out of four conditional alleles in the absence of the *Wnt1-Cre* transgene, reaching a prevalence of 83% in *Pdgfra^fl/fl^;Pdgfrb^+/fl^;Wnt1-Cre^+/+^* embryos (n = 12) (Table 2; Figure 2E,G). Further, facial hemorrhaging was noted in approximately 15% of *Pdgfra^fl/fl^;Pdgfrb^+/fl^;Wnt1-Cre^+/+^* embryos (n = 12) and double-homozygous mutant embryos (n = 14) (Table 2). These results indicate that one or both of the conditional alleles are hypomorphic. Double-homozygous mutant embryos were recovered at Mendelian frequencies at this timepoint as well (eight embryos vs. 12 expected embryos out of 93 total, χ^2^ two-tailed p = 0.2557) (Table 2). A fully-penetrant, overt facial clefting phenotype was observed in *Pdgfra^fl/fl^;Pdgfrb^+/fl^;Wnt1-Cre^+/Tg^* embryos (100%; n = 12) (Figure 2F’) and double-homozygous mutant embryos (100%; n = 8) (Figure 2H’), though not in any of the other six allele combinations from the intercrosses (n = 73) (Table 2). Facial blebbing was detected in the majority of embryos among the four genotypes containing the *Wnt1-Cre* allele and was fully penetrant in *Pdgfra^fl/fl^;Pdgfrb^+/fl^;Wnt1-Cre^+/Tg^* embryos (100%; n = 12) (Table 2; Figure 2B,D,D’,F,F’,H,H’). Similarly, facial hemorrhaging was observed in the majority of embryos containing at least three out of four conditional alleles in combination with the *Wnt1-Cre* transgene and was fully penetrant in double-homozygous mutant embryos (100%; n = 8) (Table 2; Figure 2D,D’,F,F’,H,H’). Together, these results demonstrate that *Pdgfra* and *Pdgfrb* genetically interact in the NCC lineage, with PDGFRα playing a more predominant role in NCC-mediated craniofacial development.

**Figure 2.**
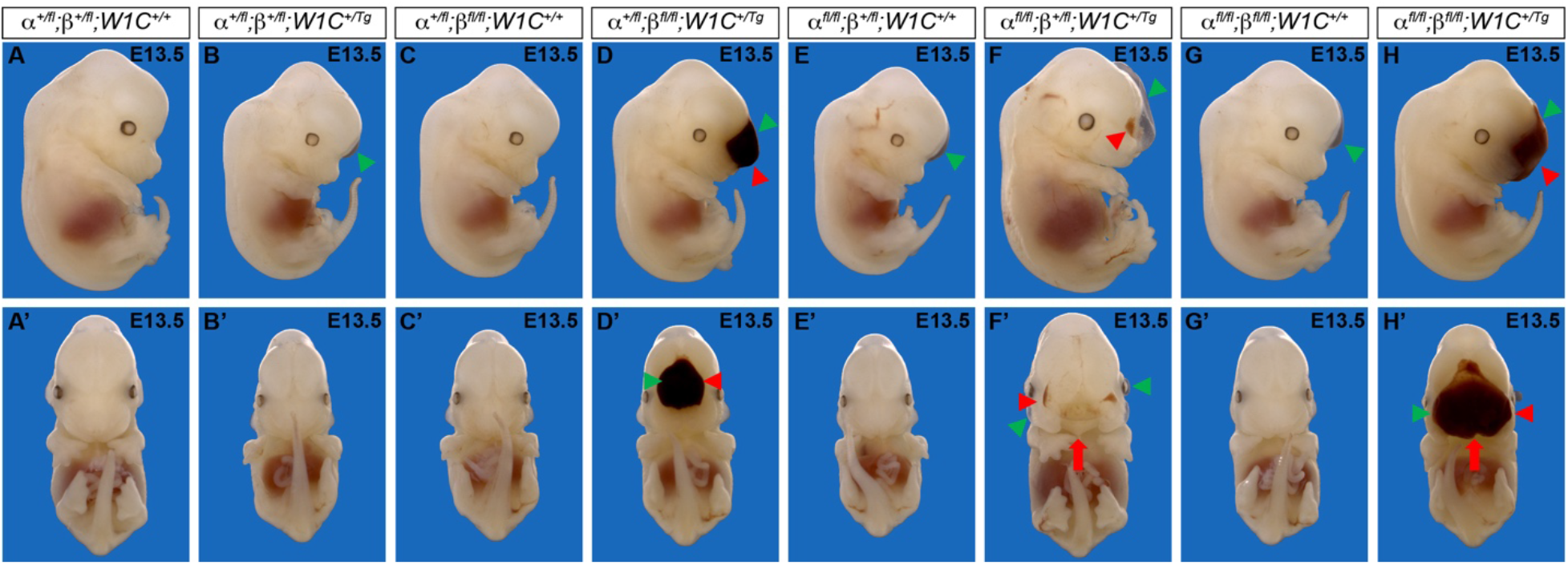
Ablation of *Pdgfra* and *Pdgfrb* in the NCC lineage results in facial clefting, blebbing and hemorrhaging at E13.5. **(A-H’)** Gross morphology of E13.5 embryos resulting from intercrosses of *Pdgfra^fl/fl^;Pdgfrb^fl/fl^* mice with *Pdgfra^+/fl^;Pdgfrb^+/fl^;Wnt1-Cre^+/Tg^* mice as viewed laterally **(A-H)** and frontally (**A’-H’)**. *Pdgfra^fl/fl^;Pdgfrb^+/fl^;Wnt1-Cre^+/Tg^* and double-homozygous mutant embryos exhibited an overt facial cleft (red arrow). Facial blebbing (green arrowheads) and facial hemorrhaging (red arrowheads) were also detected among embryos possessing a variety of allele combinations.

**Table 2.** Phenotypes of E13.5 embryos from intercrosses of *Pdgfra^fl/fl^;Pdgfrb^fl/fl^* mice with *Pdgfra^+/fl^;Pdgfrb^+/fl^;Wnt1-Cre^+/Tg^* mice.

| Genotype | Expected | Observed | Normal | Dead | Facial Cleft | Facial bleb | Facial hemorrhage |
| --- | --- | --- | --- | --- | --- | --- | --- |
| $\alpha^{+/fl}; \beta^{+/fl}; W1C^{+/+}$ | 0.125 | 0.077 | 5/5 | 3 | 0/5 | 0/5 | 0/5 |
| $\alpha^{+/fl}; \beta^{+/fl}; W1C^{+/Tg}$ | 0.125 | 0.173 | 5/16 | 2 | 0/16 | 10/16 | 2/16 |
| $\alpha^{+/fl}; \beta^{fl/fl}; W1C^{+/+}$ | 0.125 | 0.135 | 12/13 | 1 | 0/13 | 1/13 | 0/13 |
| $\alpha^{+/fl}; \beta^{fl/fl}; W1C^{+/Tg}$ | 0.125 | 0.154 | 2/13 | 3 | 0/13 | 11/13 | 9/13 |
| $\alpha^{fl/fl}; \beta^{+/fl}; W1C^{+/+}$ | 0.125 | 0.125 | 2/12 | 1 | 0/12 | 10/12 | 2/12 |
| $\alpha^{fl/fl}; \beta^{+/fl}; W1C^{+/Tg}$ | 0.125 | 0.115 | 0/12 | 0 | 12/12 | 12/12 | 9/12 |
| $\alpha^{fl/fl}; \beta^{fl/fl}; W1C^{+/+}$ | 0.125 | 0.144 | 5/14 | 1 | 0/14 | 9/14 | 2/14 |
| $\alpha^{fl/fl}; \beta^{fl/fl}; W1C^{+/Tg}$ | 0.125 | 0.077 | 0/8 | 0 | 8/8 | 7/8 | 8/8 |

### PDGFRα and, to a lesser extent, PDGFRβ regulate cNCC stream size and directional migration

We next introduced the *ROSA26^mTmG^* double-fluorescent Cre reporter allele (Muzumdar et al., 2007) into the above intercrosses to examine the timing, extent and pattern of NCC migration at E9.5-E10.5. Whereas streams entering pharyngeal arches 1 (PA1) and 2 (PA2) were readily apparent in all embryos assayed at E9.5 (Figure 3A-E”), there was a trend for the stream entering PA1 to be shorter along the anterior-posterior axis in embryos with the four experimental genotypes than in control *Pdgfra^+/+^;Pdgfrb^+/+^;Wnt1-Cre^+/Tg^* embryos (Figure 3F). Further, the anterior-posterior height and dorsal-ventral length of the stream entering PA2 were significantly shorter in *Pdgfra^fl/fl^;Pdgfrb^+/fl^;Wnt1-Cre^+/Tg^* embryos (82.22 ± 5.188 μm; 351.3 ± 13.25 μm) than in both control *Pdgfra^+/+^;Pdgfrb^+/+^;Wnt1-Cre^+/Tg^* embryos (110.0 ± 5.310 μm, p = 0.0146; 424.3 ± 14.20 μm, p = 0.0150) and *Pdgfra^+/fl^;Pdgfrb^fl/fl^;Wnt1-Cre^+/Tg^* embryos (102.5 ± 4.473 μm, p = 0.0259; 400.5 ± 12.93 μm, p = 0.0376) (Figure 3F). The height of the stream entering PA2 was also significantly shorter in double-homozygous mutant embryos (362.8 ± 11.99 μm) compared to control *Pdgfra^+/+^;Pdgfrb^+/+^;Wnt1-Cre^+/Tg^* embryos (424.3 ± 14.20 μm, p = 0.0309) (Figure 3F). These results demonstrate that combined decreases in PDGFRα and PDGFRβ signaling lead to cNCC streams entering PA1 and PA2 that are reduced in size at E9.5.

**Figure 3.**
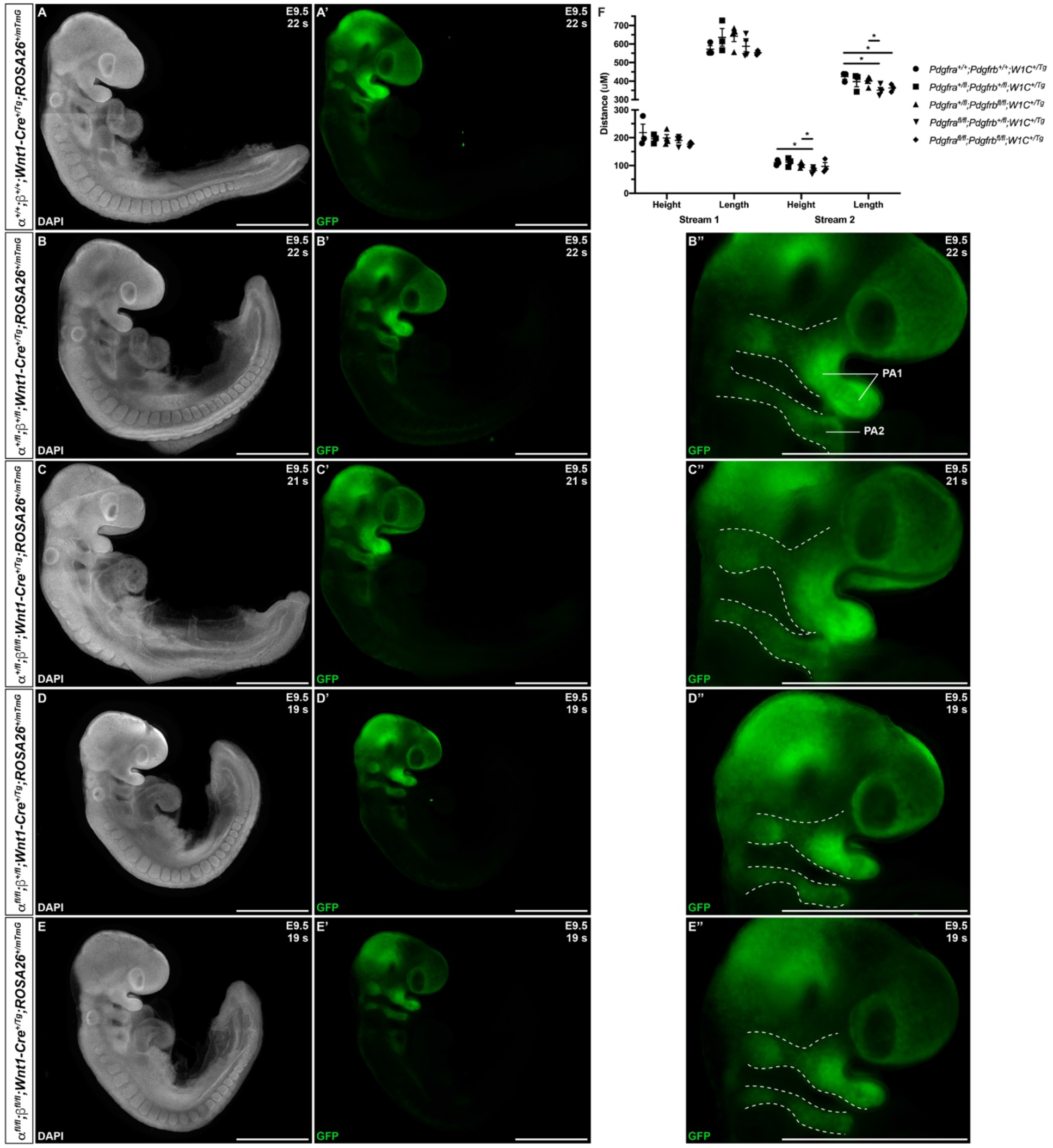
Ablation of *Pdgfra* and *Pdgfrb* in the NCC lineage leads to cNCC streams entering PA1 and PA2 that are reduced in size at E9.5. **(A-E’)** Lateral, whole-mount fluorescence images of DAPI **(A-E)** and GFP **(A’-E’)** expression across five genotypes at E9.5. **(B’’-E’’)** Zoomed-in images of GFP expression in cNCC streams (outlined by dotted lines) entering PA1 and PA2. PA1, pharyngeal arch 1; PA2, pharyngeal arch 2. **(F)** Scatter dot plot depicting the anterior-posterior heights and dorsal-ventral lengths (μm) of cNCC streams entering PA1 and PA2 across five genotypes at E9.5. Data are presented as mean ± SEM. *, p < 0.05.

At E10.5, whereas double-heterozygous mutant embryos (Figure 4B-B’”) appeared similar to control *Pdgfra^+/+^;Pdgfrb^+/+^;Wnt1-Cre^+/Tg^* embryos (Figure 4A,A’) with clearly delineated NCC streams with high cell density entering pharyngeal arches 3 (PA3) and 4 (PA4), *Pdgfra^+/fl^;Pdgfrb^fl/fl^;Wnt1-Cre^+/Tg^* embryos had streams with mild bifurcations (Figure 4C-C’”), and *Pdgfra^fl/fl^;Pdgfrb^+/fl^;Wnt1-Cre^+/Tg^* embryos had weak, diffuse streams with low cell density and more severe bifurcations (Figure 4D-D’”). Interestingly, the double-homozygous embryo phenotype was again less severe than that of *Pdgfra^fl/fl^;Pdgfrb^+/fl^;Wnt1-Cre^+/Tg^* embryos. Double-homozygous mutant embryos also exhibited weak streams with low cell density, but no apparent diffusion of the streams, and only mild bifurcations (Figure 4E’-E’”). While the anterior-posterior heights of the streams entering PA3 and PA4 did not vary significantly among control *Pdgfra^+/+^;Pdgfrb^+/+^;Wnt1-Cre^+/Tg^* embryos and embryos with the four experimental genotypes, there was a trend for the streams from double-heterozygous mutant embryos and *Pdgfra^fl/fl^;Pdgfrb^+/fl^;Wnt1-Cre^+/Tg^* embryos to be taller than those from *Pdgfra^+/fl^;Pdgfrb^fl/fl^;Wnt1-Cre^+/Tg^* embryos and double-homozygous mutant embryos (Figure 4F). Alternatively, while the dorsal-ventral lengths of the streams entering PA3 and PA4 also did not vary significantly between genotypes, there was a trend for the streams from *Pdgfra^fl/fl^;Pdgfrb^+/fl^;Wnt1-Cre^+/Tg^* embryos and double-homozygous mutant embryos to be longer than those from double-heterozygous mutant embryos and *Pdgfra^+/fl^;Pdgfrb^fl/fl^;Wnt1-Cre^+/Tg^* embryos (Figure 4F).

**Figure 4.**
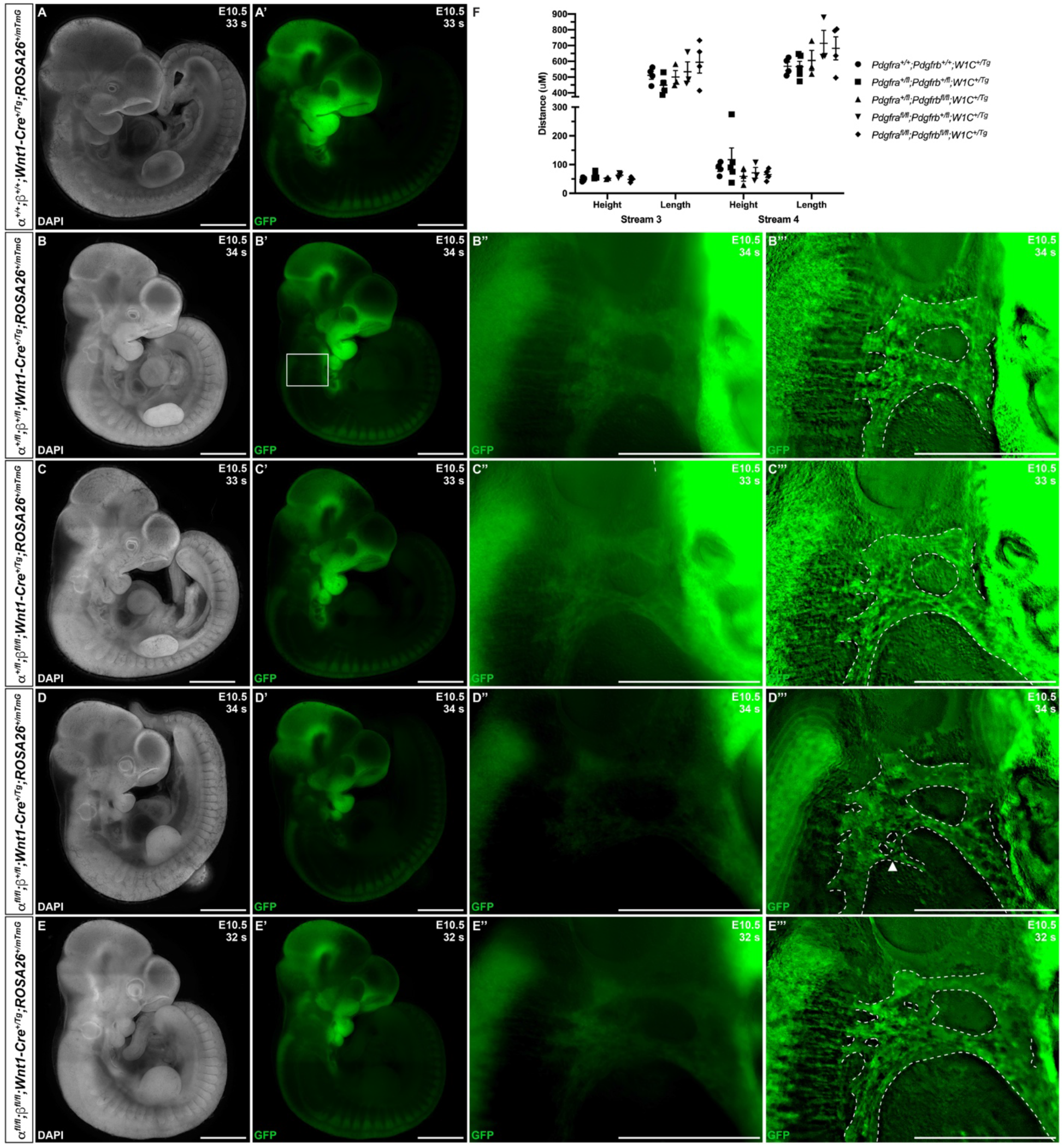
Ablation of *Pdgfra* and *Pdgfrb* in the NCC lineage results in longer, more diffuse cNCC streams along the dorsal-ventral axis entering PA3 and PA4 at E10.5, with increased incidences of stream bifurcations and intermingling. **(A-E’)** Lateral, whole-mount fluorescence images of DAPI **(A-E)** and GFP **(A’-E’)** expression across five genotypes at E10.5. **(B’’-E’”)** Zoomed-in images of GFP expression in cNCC streams (outlined by dotted lines) entering PA3 and PA4. Arrowhead indicates an example of a bifurcated cNCC stream. **(F)** Scatter dot plot depicting the anterior-posterior heights and dorsal-ventral lengths (μm) of cNCC streams entering PA3 and PA4 across five genotypes at E10.5. Data are presented as mean ± SEM.

The above E10.5 embryos were then scored for bifurcations in streams entering PA3-PA4 and intermingling of the two streams. For a handful of embryos with a relatively high number of somite pairs (≥39), the stream entering PA3 was no longer visible and hence was not assayed for bifurcation or intermingling with the stream entering PA4. The stream entering PA3 was not bifurcated in any of the double-heterozygous mutant embryos (n = 4), but was found to be bifurcated in 33% of *Pdgfra^+/fl^;Pdgfrb^fl/fl^;Wnt1-Cre^+/Tg^* embryos (n = 3), 50% of *Pdgfra^fl/fl^;Pdgfrb^+/fl^;Wnt1-Cre^+/Tg^* embryos (n = 2) and 67% of double-homozygous mutant embryos (n= 3) (Table 3). Bifurcation of the stream entering PA4 was observed in 40% of double-heterozygous mutant embryos (n = 5), 67% of *Pdgfra^fl/fl^;Pdgfrb^+/fl^;Wnt1-Cre^+/Tg^* embryos (n = 3) and was fully penetrant in *Pdgfra^+/fl^;Pdgfrb^fl/fl^;Wnt1-Cre^+/Tg^* embryos (100%; n= 3) and double-homozygous mutant embryos (100%; n = 4) (Table 3). Finally, the streams entering PA3-PA4 were intermingled in 75% of double-heterozygous mutant embryos (n = 4) and in all *Pdgfra^+/fl^;Pdgfrb^fl/fl^;Wnt1-Cre^+/Tg^* embryos (100%; n = 3), *Pdgfra^fl/fl^;Pdgfrb^+/fl^;Wnt1-Cre^+/Tg^* embryos (100%; n = 2) and double-homozygous mutant embryos (100%; n = 3) (Table 3). Taken together, the results at E10.5 indicate that combined decreases in PDGFRα and PDGFRβ signaling lead to longer, more diffuse cNCC streams along the dorsal-ventral axis entering PA3 and PA4, with increased incidences of stream bifurcations and intermingling, perhaps indicative of defects in NCC directional migration.

**Table 3.** Bifurcation and intermingling of NCC streams entering PA3 and PA4 at E 10.5.

| Genotype | Bifurcated Stream 3 | Bifurcated Stream 4 | Intermingling of Streams 3 and 4 |
| --- | --- | --- | --- |
| $\alpha^{+/fl}; \beta^{+/fl}; W1C^{+/Tg}$ | 0/4 | 2/5 | 3/4 |
| $\alpha^{+/fl}; \beta^{fl/fl}; W1C^{+/Tg}$ | 1/3 | 3/3 | 3/3 |
| $\alpha^{fl/fl}; \beta^{+/fl}; W1C^{+/Tg}$ | 1/2 | 2/3 | 2/2 |
| $\alpha^{fl/fl}; \beta^{fl/fl}; W1C^{+/Tg}$ | 2/3 | 4/4 | 3/3 |

Finally, to assess the extent of NCCs and their derivatives in the facial processes at E9.5 and E10.5, we quantified GFP expression in frontal views of the head in control *Pdgfra^+/+^;Pdgfrb^+/+^;Wnt1-Cre^+/Tg^* embryos and among embryos with the four experimental genotypes. At E9.5, there were noticeable decreases in GFP intensity in the facial processes of experimental embryos (Figure 5B’-E’) compared to control embryos (Figure 5A’), particularly in the frontonasal and maxillary prominences. GFP fluorescence values were significantly decreased in *Pdgfra^fl/fl^;Pdgfrb^+/fl^;Wnt1-Cre^+/Tg^* embryos (8.449 x 10^8^ ± 7.256 x 10^7^) compared to control *Pdgfra^+/+^;Pdgfrb^+/+^;Wnt1-Cre^+/Tg^* embryos (2.079 x 10^9^ ± 2.539 x 10^8^) and double-heterozygous mutant embryos (1.373 x 10^9^ ± 1.283 x 10^8^). Moreover, while double-homozygous mutant embryos had higher GFP fluorescence values than *Pdgfra^fl/fl^;Pdgfrb^+/fl^;Wnt1-Cre^+/Tg^* embryos, GFP fluorescence was significantly decreased in double-homozygous mutant embryos (1.088 x 10^9^ ±1.022 x 10^8^) compared to control *Pdgfra^+/+^;Pdgfrb^+/+^;Wnt1-Cre^+/Tg^* embryos (2.079 x 10^9^ ± 2.539 x 10^8^). At E10.5, there was a marked decrease in GFP intensity in the facial processes of double-heterozygous mutant embryos (Figure 5H’) compared to control *Pdgfra^+/+^;Pdgfrb^+/+^;Wnt1-Cre^+/Tg^* embryos (Figure 5G’) and a further decrease in *Pdgfra^+/fl^;Pdgfrb^fl/fl^;Wnt1-Cre^+/Tg^* embryos (Figure 5I’), *Pdgfra^fl/fl^;Pdgfrb^+/fl^;Wnt1-Cre^+/Tg^* embryos (Figure 5j’) and double-homozygous mutant embryos (Figure 5K’). Not surprisingly, GFP fluorescence values increased with the number of somite pairs, as NCC progenitors proliferate and differentiate over time (Figure 5L). However, for embryos with 31-35 somite pairs, relative fluorescence units decreased as additional alleles were ablated, with *Pdgfra^fl/fl^;Pdgfrb^+/fl^;Wnt1-Cre^+/Tg^* and double-homozygous mutant embryos having the lowest, and essentially equal, GFP fluorescence values (Figure 5L). Collectively, our assessment of cNCC migration in the context of *Pdgfra* and *Pdgfrb* ablation demonstrates that signaling through these receptors contributes to several aspects of NCC activity, including stream size, directional migration and, ultimately, the extent of their derivatives in the facial prominences. Importantly, PDGFRα signaling appears to play a more predominant role in cNCC migration than PDGFRβ.

**Figure 5.**
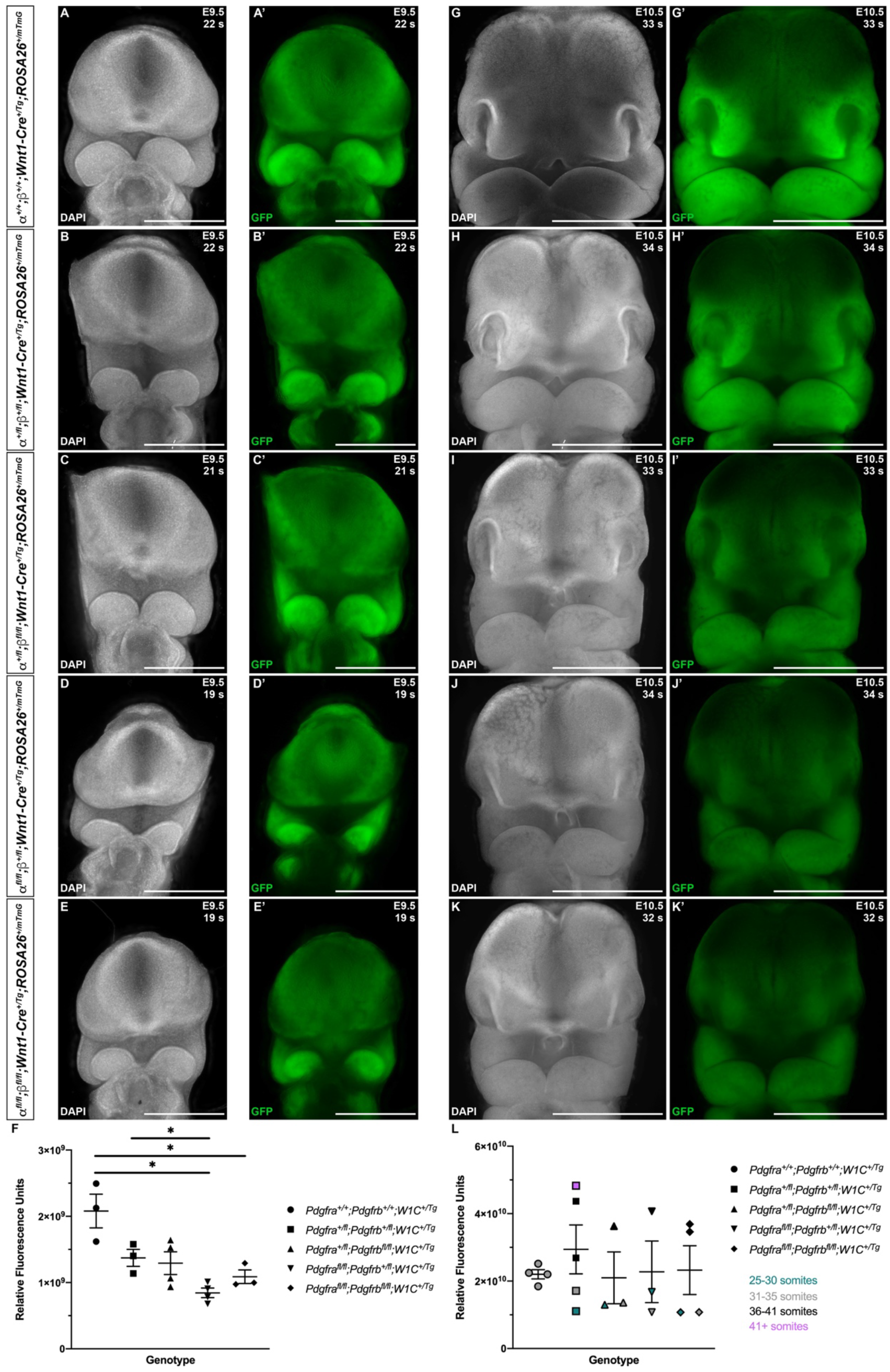
Ablation of *Pdgfra* and *Pdgfrb* in the NCC lineage leads to decreased NCC derivatives in the facial prominences at mid-gestation. **(A-K’)** Frontal, whole-mount fluorescence images of DAPI **(A-E, G-K)** and GFP **(A’-E’, G’-K’)** expression across five genotypes at E9.5 **(A-E’)** and E10.5 **(G-K’)**. **(F)** Scatter dot plot depicting GFP fluorescence intensity across five genotypes at E9.5. Data are presented as mean ± SEM. *, p < 0.05. **(L)** Scatter dot plot depicting GFP fluorescence intensity across five genotypes at E10.5. Data are presented as mean ± SEM. Colors correspond to number of somite pairs in assayed embryos.

### PDGFRβ plays a more dominant role in proliferation of the facial mesenchyme than PDGFRα

We next examined levels of cell death amongst one control, *Pdgfra^+/fl^;Pdgfrb^+/fl^;Wnt1-Cre*^+/+^, and the four experimental genotypes containing the *Wnt1-Cre* transgene via terminal deoxynucleotidyl transferase-mediated dUTP nick end labeling (TUNEL). At E10.5, the percentage of TUNEL-positive cells was determined within the mesenchyme of the lateral and medial nasal processes, as well as the maxillary and mandibular prominences. The percentage of TUNEL-positive cells was higher in the medial nasal processes than the other locations at this timepoint for all genotypes (Figure 6A). Further, all experimental genotypes had a non-statistically significant decrease in the percentage of TUNEL-positive cells compared to the control genotype in both the lateral and medial nasal processes (Figure 6A). While the level of cell death did not vary significantly between the five genotypes within the maxillary and mandibular prominences, *Pdgfra^fl/fl^;Pdgfrb^+/fl^;Wnt1-Cre^+/Tg^* embryos and double-heterozygous mutant embryos had the highest and second-highest percentages of TUNEL-positive cells, respectively, at these locations (Figure 6A). In the mandibular prominence, there was a trend for each of the experimental genotypes to have a higher percentage of TUNEL-positive cells when compared to control embryos (Figure 6A). At E13.5, the percentage of TUNEL-positive cells was determined within the mesenchyme of the nasal septum and anterior, middle and posterior secondary palatal shelves. The percentage of TUNEL-positive cells was higher in the nasal septum than in the secondary palatal shelves for all genotypes, consistent with the relatively high level of TUNEL-positive cells in the medial nasal processes three days earlier at E10.5. Two genotypes, *Pdgfra^+/fl^;Pdgfrb^fl/fl^;Wnt1-Cre^+/Tg^* embryos and *Pdgfra^fl/fl^;Pdgfrb^+/fl^;Wnt1-Cre^+/Tg^* embryos, had a non-statistically significant increase in the percentage of TUNEL-positive cells compared to the control genotype at this location (Figure 6B). While the level of cell death did not vary significantly between the five genotypes within the secondary palatal shelves, there was a trend for each of the experimental genotypes to have a lower percentage of TUNEL-positive cells in the anterior palatal shelves when compared to control embryos (Figure 6B). In the middle palatal shelves, three genotypes, double-heterozygous mutant embryos, *Pdgfra^+/fl^;Pdgfrb^fl/fl^;Wnt1-Cre^+/Tg^* embryos and *Pdgfra^fl/fl^;Pdgfrb^+/fl^;Wnt1-Cre^+/Tg^* embryos, had a non-statistically significant increase in the percentage of TUNEL-positive cells compared to the control genotype (Figure 6B). Similarly, in the posterior palatal shelves, three genotypes, double-heterozygous mutant embryos, *Pdgfra^fl/fl^;Pdgfrb^+/fl^;Wnt1-Cre^+/Tg^* embryos and double-homozygous mutant embryos, had a non-statistically significant increase in the percentage of TUNEL-positive cells compared to the control genotype (Figure 6B). Despite these modest trends, the combined TUNEL assay results demonstrate that neither PDGFRα nor PDGFRβ signaling plays a critical role in cNCC-derived facial mesenchyme survival during mid-gestation.

**Figure 6.**
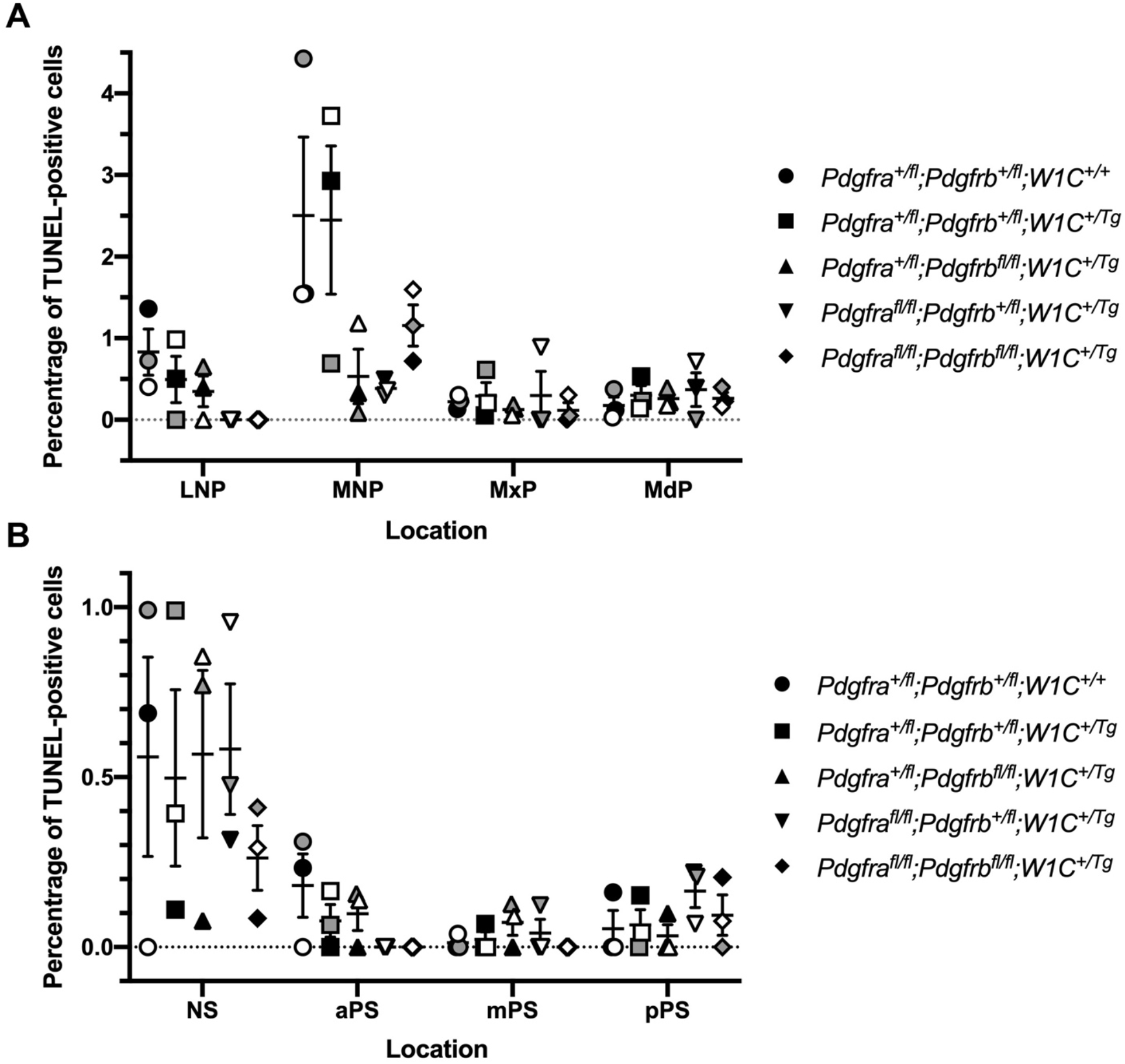
Neither PDGFRα nor PDGFRβ signaling plays a critical role in cNCC-derived facial mesenchyme survival during mid-gestation. **(A)** Scatter dot plot depicting the percentage of TUNEL-positive cells in the nasal processes and facial prominences across five genotypes at E10.5. Data are presented as mean ± SEM. Shades correspond to independent experiments across three biological replicates. LNP, lateral nasal process; MNP, medial nasal process; MxP, maxillary prominence; MdP, mandibular prominence. **(B)** Scatter dot plot depicting the percentage of TUNEL-positive cells in the nasal septum and secondary palatal shelves across five genotypes at E13.5. Data are presented as mean ± SEM. Shades correspond to independent experiments across three biological replicates. NS, nasal septum; aPS, anterior secondary palatal shelves; mPS, middle secondary palatal shelves; pPS, posterior secondary palatal shelves.

We similarly examined levels of cell proliferation amongst one control, *Pdgfra^+/fl^;Pdgfrb^+/fl^;Wnt1-Cre*^+/+^, and the four experimental genotypes containing the *Wnt1-Cre* transgene via Ki67 immunofluorescence analysis. At E10.5, the percentage of Ki67-positive cells was determined within the mesenchyme of the lateral and medial nasal processes, as well as the maxillary and mandibular prominences. The percentage of Ki67-positive cells was highest in the lateral nasal processes and lowest in the mandibular prominence for all genotypes (Figure 7A). The level of cell proliferation did not vary significantly between the five genotypes at any of the locations examined, with the exception of a significant increase in the percentage of Ki67-positive cells in the lateral nasal processes of double-homozygous mutant embryos (4.704 ± 0.4459) compared to *Pdgfra^fl/fl^;Pdgfrb^+/fl^;Wnt1-Cre^+/Tg^* embryos (3.004 ± 0.3356, p = 0.0420) (Figure 7A). All experimental genotypes had a non-statistically significant decrease in the percentage of Ki67-positive cells compared to the control genotype in the maxillary prominences (Figure 7A). Interestingly, the percentage of Ki67-positive cells was consistently lower in *Pdgfra^+/fl^;Pdgfrb^fl/fl^;Wnt1-Cre^+/Tg^* embryos and *Pdgfra^fl/fl^;Pdgfrb^+/fl^;Wnt1-Cre^+/Tg^* embryos than double-homozygous mutant embryos at all locations at this timepoint (Figure 7A). As above with the TUNEL analysis, at E13.5, the percentage of Ki67-positive cells was determined within the mesenchyme of the nasal septum and anterior, middle and posterior secondary palatal shelves. The percentage of Ki67-positive cells was consistently lower in the nasal septum than the secondary palatal shelves (Figure 7B). While the level of proliferation did not vary significantly between the five genotypes in the nasal septum and along the anterior-posterior axis of the secondary palatal shelves, there were trends for each of the experimental genotypes to have a lower percentage of Ki67-positive cells in the nasal septum and middle palatal shelves and a higher percentage of Ki67-positive cells in the posterior palatal shelves when compared to these same locations in control embryos (Figure 7B). Intriguingly, *Pdgfra^+/fl^;Pdgfrb^fl/fl^;Wnt1-Cre^+/Tg^* embryos had a consistently lower percentage of Ki67-positive cells in the nasal septum and throughout the secondary palatal shelves than *Pdgfra^fl/fl^;Pdgfrb^+/fl^;Wnt1-Cre^+/Tg^* embryos (Figure 7B). These findings indicate that both PDGFRα and PDGFRβ promote cell proliferation in the craniofacial mesenchyme, with PDGFRβ potentially playing a more predominant role in this context.

**Figure 7.**
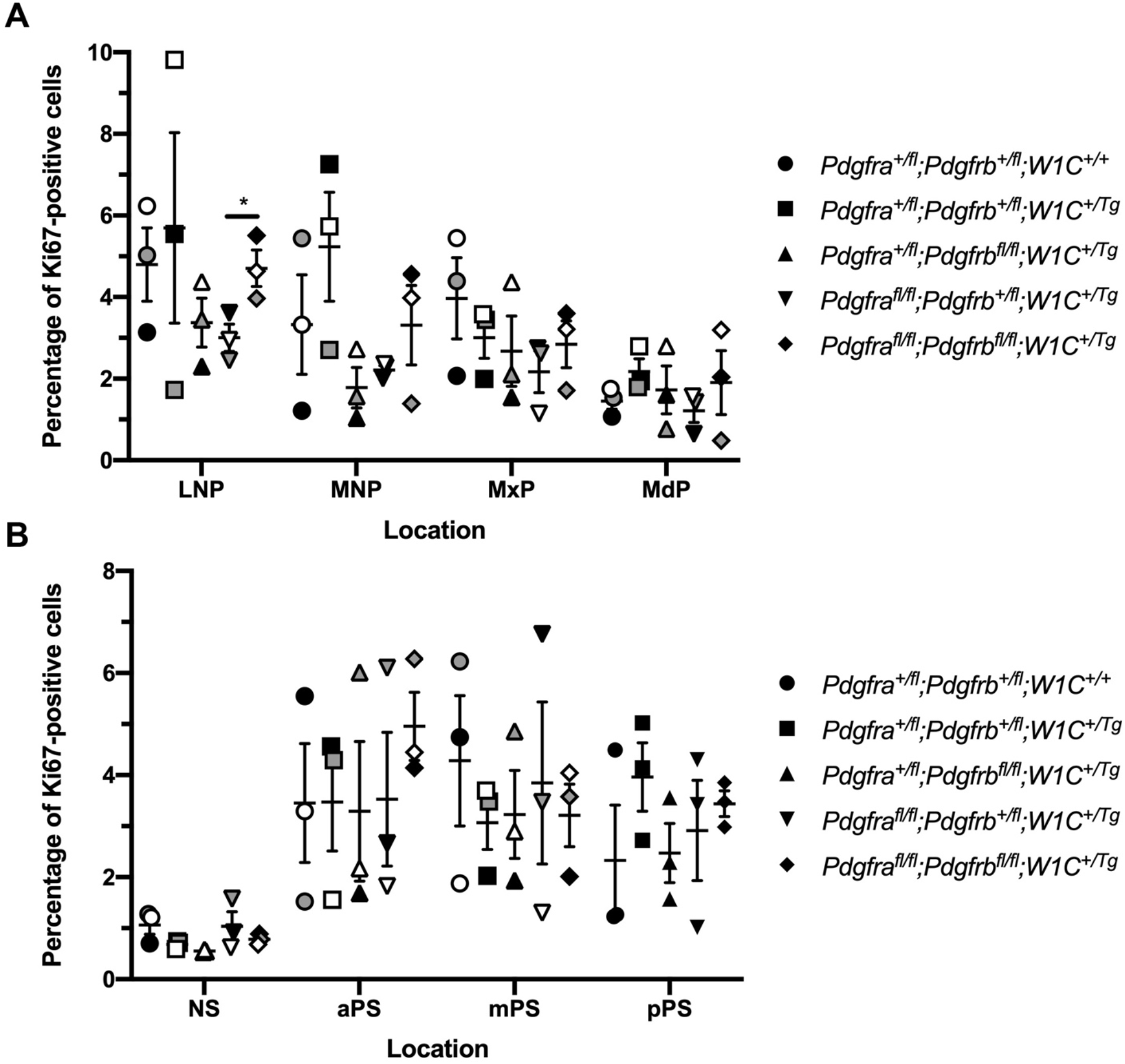
PDGFRβ plays a more dominant role in proliferation of the craniofacial mesenchyme than PDGFRα during mid-gestation. **(A)** Scatter dot plot depicting the percentage of Ki67-positive cells in the nasal processes and facial prominences across five genotypes at E10.5. Data are presented as mean ± SEM. *, p < 0.05. Shades correspond to independent experiments across three biological replicates. LNP, lateral nasal process; MNP, medial nasal process; MxP, maxillary prominence; MdP, mandibular prominence. **(B)** Scatter dot plot depicting the percentage of Ki67-positive cells in the nasal septum and secondary palatal shelves across five genotypes at E13.5. Data are presented as mean ± SEM. Shades correspond to independent experiments across three biological replicates. NS, nasal septum; aPS, anterior secondary palatal shelves; mPS, middle secondary palatal shelves; pPS, posterior secondary palatal shelves.

We subsequently sought to determine the individual contribution of PDGFRα and PDGFRβ to proliferation of the craniofacial mesenchyme and to distinguish any potential proliferation defects from more wide-spread phenotypes observed upon ablation of *Pdgfra* or *Pdgfrb* in the NCC lineage. To do this, primary MEPM cells were dissected from E13.5 control (*Pdgfra^+/fl^;Wnt1-Cre^+/Tg^* or *Pdgfrb^+/fl^;Wnt1-Cre^+/Tg^*) and conditional knock-out (*Pdgfra^fl/fl^;Wnt1-Cre^+/Tg^* or *Pdgfrb^fl/fl^;Wnt1-Cre^+/Tg^*) littermate embryos for use in cell growth assays (Figure 8A). Primary MEPM cells are a faithful surrogate for embryonic facial mesenchyme, as wild-type cells uniformly express both PDGFRα and PDGFRβ, as well as numerous additional markers of palatal mesenchyme cells *in vivo*, and are responsive to stimulation with PDGF-AA, PDGF-BB and PDGF-DD ligand (He and Soriano, 2013; Fantauzzo and Soriano, 2014, 2016, 2017; Vasudevan and Soriano, 2014; Vasudevan et al., 2015). Even after a single day in growth medium containing 10% FBS, control *Pdgfrb^+/fl^;Wnt1-Cre^+/Tg^* cells (0.1077 ± 0.01233 arbitrary units (AU)) (Figure 8C) had grown about half as much as control *Pdgfra^+/fl^;Wnt1-Cre^+/Tg^* cells (0.2217 ± 0.07322 AU) (Figure 8B). All cells grown in starvation medium containing 0.1% FBS, both control and conditional knock-out, immediately proliferated less than cells of the same genotypes grown in growth medium (Figure 8B,C). Importantly, conditional knock-out cells consistently fared worse than their control counterparts in both growth and starvation medium, though this difference was more pronounced in *Pdgfrb^+/fl^;Wnt1-Cre^+/Tg^* versus *Pdgfrb^fl/fl^;Wnt1-Cre^+/Tg^* cells following six days in culture. At this time, control *Pdgfra^+/fl^;Wnt1-Cre^+/Tg^* cells cultured in growth medium (0.8773 ± 0.08867 AU) had proliferated approximately 1.8 times the extent of *Pdgfra^fl/fl^;Wnt1-Cre^+/Tg^* cells (0.4885 ± 0.03203 AU, p = 0.0357) (Figure 8B), while control *Pdgfrb^+/fl^;Wnt1-Cre^+/Tg^* cells (0.5897 ± 0.03588 AU) cultured in growth medium had an absorbance reading roughly 2.5 times that of *Pdgfrb^fl/fl^;Wnt1-Cre^+/Tg^* cells (0.2394 ± 0.05482 AU, p = 0.0018) (Figure 8C). Similarly, control *Pdgfra^+/fl^;Wnt1-Cre^+/Tg^* cells cultured in starvation medium (0.2953 ± 0.06842 AU) had proliferated approximately 1.5 times the extent of *Pdgfra^fl/fl^;Wnt1-Cre^+/Tg^* cells (0.2013 ± 0.01605 AU, p = 0.3012) (Figure 8B), while control *Pdgfrb^+/fl^;Wnt1-Cre^+/Tg^* cells (0.2047 ± 0.009821 AU) cultured in growth medium had an absorbance reading roughly 1.9 times that of *Pdgfrb^fl/fl^;Wnt1-Cre^+/T^g* cells (0.1084 ± 0.01588 AU, p = 0.0022) (Figure 8C). Further, none of the PDGF-AA, PDGF-BB nor PDGF-DD ligand treatments led to significantly more growth than that observed in the absence of ligand for cells of all genotypes cultured in both growth and starvation medium (Figure 8B,C), with the exception of control *Pdgfrb^+/fl^;Wnt1-Cre^+/Tg^* cells cultured in starvation medium in the absence (0.2047 ± 0.009821 AU) or presence of PDGF-AA ligand treatment (0.1580 ± 0.01286 AU, p = 0.0486) (Figure 8C). Interestingly, PDGF-AA ligand treatment, which has been shown to exclusively activate PDGFRα homodimer signaling in this context and not PDGFRα/β heterodimer nor PDGFRβ homodimer signaling (Fantauzzo and Soriano, 2017), consistently resulted in less cell growth than PDGF-BB and/or PDGF-DD ligand treatments, though these differences were not statistically significant (Figure 8B,C). Finally, whereas control *Pdgfra^+/fl^;Wnt1-Cre^+/Tg^* cells versus *Pdgfra^fl/fl^;Wnt1-Cre^+/Tg^* cells did not exhibit a significant difference in proliferation upon ligand treatment when cultured in growth or starvation medium, control *Pdgfrb^+/fl^;Wnt1-Cre^+/Tg^* cells significantly out-performed *Pdgfrb^fl/fl^;Wnt1-Cre^+/Tg^* cells when cultured in growth medium in response to PDGF-AA ligand treatment (0.5167 ± 0.01913 versus 0.2912 ± 0.07210, p = 0.0332) and when cultured in starvation medium in response to PDGF-DD ligand treatment (0.1770 ± 0.005859 versus 0.1196 ± 0.02084, p = 0.0493) (Figure 8B,C). Taken together, these results confirm the Ki67 immunofluorescence analyses above and reveal that PDGFRβ plays a more dominant role in proliferation of the facial mesenchyme than PDGFRα.

**Figure 8.**
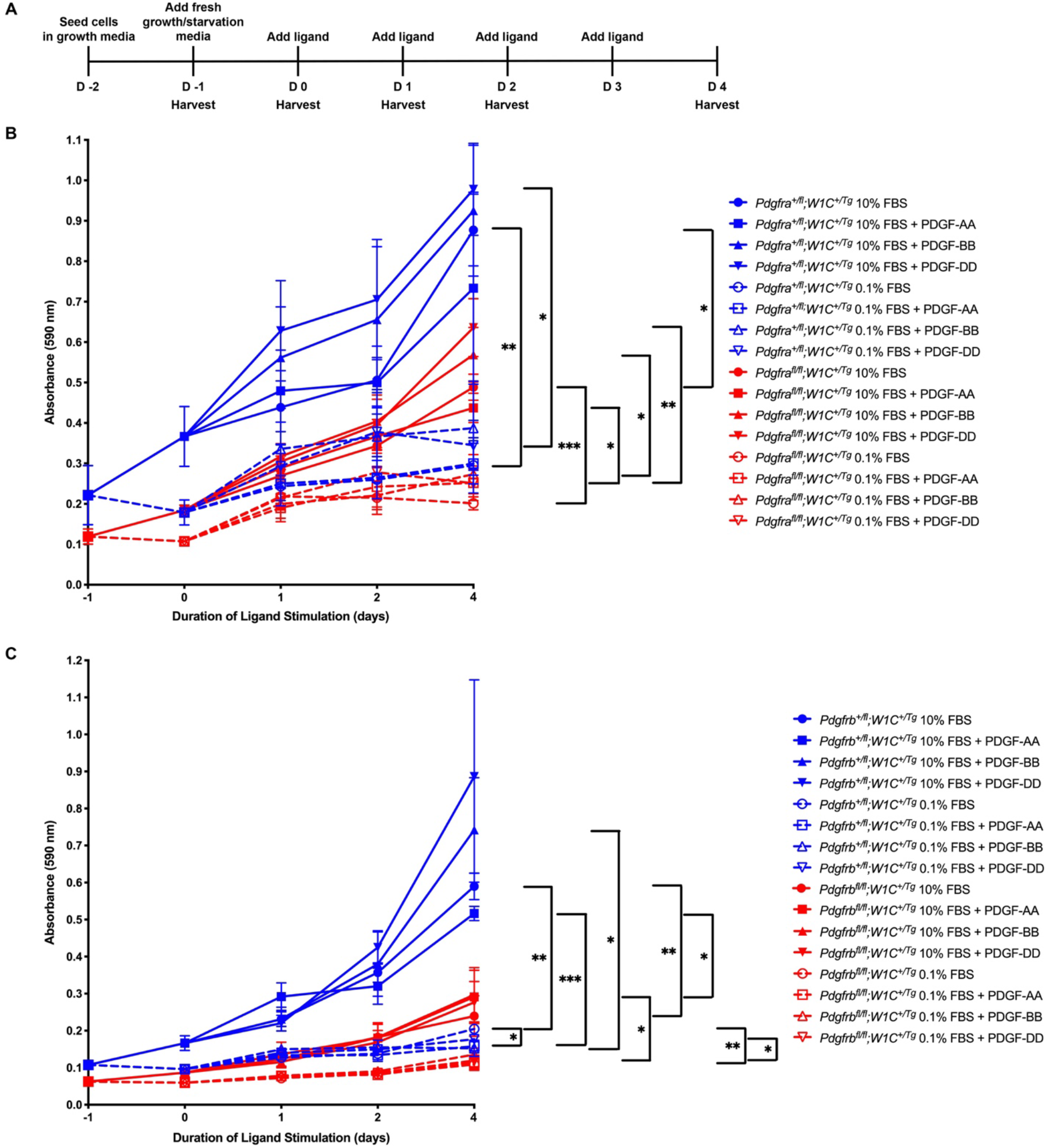
PDGFRβ plays a more dominant role in proliferation of primary MEPM cells than PDGFRα. **(A)** Experimental design for cell growth assays. **(B)** Line graph depicting absorbance values at 590 nm in *Pdgfra^+/fl^;Wnt1-Cre^+/Tg^* versus *Pdgfra^fl/fl^;Wnt1-Cre^+/Tg^* primary MEPM cells across conditions. Data are presented as mean ± SEM. *, p < 0.05; **, p < 0.01; ***, p < 0.001. **(C)** Line graph depicting absorbance values at 590 nm in *Pdgfrb^+/fl^;Wnt1-Cre^+/Tg^* versus *Pdgfrb^fl/fl^;Wnt1-Cre^+/Tg^* primary MEPM cells across conditions. Data are presented as mean ± SEM. *, p < 0.05; **, p < 0.01; ***, p < 0.001.

## Discussion

Here we report the first detailed phenotypic characterization of mouse embryos in which both *Pdgfra* and *Pdgfrb* are ablated in the NCC lineage. Our results reveal that the two receptors genetically interact in this lineage during embryogenesis, as phenotypes observed in an allelic series of mutant embryos often worsened with the addition of conditional alleles. We characterized defects in craniofacial development at mid-gestation resulting from combined loss of *Pdgfra* and *Pdgfrb*, including incidences of facial clefting, blebbing and hemorrhaging. These results confirm the phenotypes we observed from mid-to-late-gestation upon combining the constitutive *Pdgfra^PI3K^* allele together with the conditional *Pdgfrb^fl^* allele and the *Wnt1-Cre* driver (Fantauzzo and Soriano, 2016) and significantly extend those findings by exploring the cellular mechanisms through which these phenotypes arise. The defects observed here were shown to stem from decreased cNCC stream size and aberrant cNCC directional migration, as well as reduced proliferation of the facial mesenchyme upon combined decreases in PDGFRα and PDGFRβ signaling. These findings are the first to demonstrate a role for PDGFRβ in regulating each of these processes in the developing mouse embryo. Importantly, we found that PDGFRα plays a predominant role in cNCC migration while PDGFRβ primarily contributes to proliferation of the facial mesenchyme.

Our E13.5 gross morphology results indicate that one or both of the conditional alleles used in this study are hypomorphic, as facial blebbing and facial hemorrhaging were detected in a subset of embryos upon combination of at least 3 out of 4 conditional alleles in the absence of the *Wnt1-Cre* transgene. While mice heterozygous for a *Pdgfra* null allele are viable (Soriano, 1997), *Pdgfra^fl/-^* embryos are not, exhibiting multiple phenotypes such as spina bifida and cleft palate (Tallquist and Soriano, 2003; McCarthy et al., 2016). Further, *Pdgfra^fl/fl^* mice in our own colony, which are maintained through homozygous intercrosses, generate small litters (average litter size of 4.2 pups at 5-10 days after birth compared to an average of 5.8 pups for wild-type 129S4 litters; p = 0.0013) and have shortened snouts with a pigment defect at the facial midline (data not shown). It has been hypothesized that these hypomorphic phenotypes arise due to the presence of a neomycin resistance cassette in the floxed allele that reduces expression of *Pdgfra* (Tallquist and Soriano, 2003). Hypomorphic phenotypes have not previously been attributed to the *Pdgfrb^fl^* allele, and *Pdgfrb^fl/fl^* mice in our colony, which are also maintained through homozygous intercrosses, give birth to litters of expected sizes (average litter size of 6.2 pups at 5-10 days after birth compared to an average of 5.8 pups for wild-type 129S4 litters; p = 0.2998).

Interestingly, in several parameters examined here, including distance between the nasal pits at E10.5, heights and lengths of cNCC streams entering PA2, GFP intensity in the facial processes at E9.5 and the percentage of Ki67-positive cells in the lateral nasal processes at E10.5, the phenotype of *Pdgfra^fl/fl^;Pdgfrb^+/fl^;Wnt1-Cre^+/Tg^* embryos was more severe than that of double-homozygous mutant embryos. This result is contrary to our previous observations in which *Pdgfra^PI3K/PI3K^;Pdgfrb^+/fl^;Wnt1-Cre^+/Tg^* embryos did not exhibit facial clefting at E13.5, while this phenotype was fully penetrant in *Pdgfra^PI3K/PI3K^;Pdgfrb^fl/fl^;Wnt1-Cre^+/Tg^* embryos (Fantauzzo and Soriano, 2016). The most likely explanation for this finding is that reduced, but not absent, PDGFRβ signaling has a negative effect on cNCC activity and subsequent facial development in a context in which PDGFRα signaling is completely abolished, as observed here. Further studies will be required to determine the mechanism(s) by which this phenomenon occurs.

In *Xenopus, pdgfra* is expressed by pre-migratory and migratory cNCCs, while its ligand *pdgfa* is expressed in pre-migratory NCCs and the tissues surrounding migratory NCCs (Bahm et al., 2017). Functional studies revealed dual roles for PDGF-A-dependent PDGFRα signaling in NCC development. During early NCC migration, PI3K/Akt-mediated PDGFRα signaling cell autonomously upregulates N-cadherin to promote contact inhibition of locomotion and cell dispersion. Following initiation of the epithelial-to-mesenchymal transition, migrating NCCs chemotax towards PDGF-A ligand in the surrounding tissue, resulting in directional migration (Bahm et al., 2017). The ligand *pdgfb* is also expressed in tissues adjacent to migrating NCCs in *Xenopus* embryos (Giannetti et al., 2016) and knock-down of this ligand results in impaired cNCC migration and defective development of the craniofacial cartilages and cranial nerves in a subset of morpholino-injected embryos (Corsinovi et al., 2019). In zebrafish, *pdgfra* is similarly expressed by pre-migratory and migratory cNCCs, while its ligand *pdgfaa* is correspondingly expressed at early stages in the midbrain and later in the oral ectoderm (Eberhart et al., 2008). A hypomorphic zebrafish mutant of *pdgfra* exhibits palatal clefting and a shortened neurocrania due to defective cNCC migration (Eberhart et al., 2008; McCarthy et al., 2016). *Pdgfrb* is also expressed by migratory cNCCs in zebrafish and the phenotypes observed in *pdgfra* mutants are exacerbated in double *pdgfra;pdgfrb* mutant fish in which cNCCs fail to properly condense in the maxillary domain (McCarthy et al., 2016). In contrast to a previous study in which cNCC migration was reportedly unperturbed upon combined ablation of *Pdgfra* and *Pdgfrb* in the murine NCC lineage (Richarte et al., 2007), our results confirm the findings in lower vertebrates that both receptors play a role in NCC migration and that aspects of the phenotype observed upon conditional ablation of *Pdgfra* in the NCC lineage are exacerbated in double-homozygous mutant embryos.

In summary, our findings provide insight into the distinct mechanisms by which PDGFRα and PDGFRβ signaling regulate cNCC activity and subsequent craniofacial development in a mammalian system. Future studies will seek to identify the intracellular signaling molecules and gene expression responses that mediate the effects of these receptors on migration and proliferation.

## Author Contributions

KF conceived and designed the study. JM, RL and KF performed experimentation. JM and KF analyzed data. KF wrote the original draft of the manuscript, which was revised and edited in an iterative process with JM.

## Acknowledgements

We are grateful to Damian Garno and Elliott Brooks for technical assistance. We thank members of the Fantauzzo laboratory for their helpful discussions and critical comments on the manuscript.

